# RqcH supports survival in the absence of non-stop ribosome rescue factors

**DOI:** 10.1101/2024.07.12.603306

**Authors:** Katrina Callan, Cassidy R. Prince, Heather A. Feaga

**Affiliations:** Department of Microbiology, Cornell University, Ithaca, NY 14853

**Keywords:** ribosome rescue, protein synthesis, translation, rqcH, nonstop mRNA

## Abstract

Ribosomes frequently translate truncated or damaged mRNAs due to the extremely short half-life of mRNAs in bacteria. When ribosomes translate mRNA that lacks a stop codon (non-stop mRNA), specialized pathways are required to rescue the ribosome from the 3’ end of the mRNA. The most highly conserved non-stop rescue pathway is *trans*-translation, which is found in greater than 95% of bacterial genomes. In all Proteobacteria that have been studied, the alternative non-stop ribosome rescue factors, ArfA and ArfB, are essential in the absence of *trans*-translation. Here, we investigate the interaction between non-stop rescue pathways and RqcH, a ribosome quality control factor that is broadly conserved outside of Proteobacteria. RqcH does not act directly on non-stop ribosomes but adds a degron tag to stalled peptides that obstruct the large ribosomal subunit, which allows the stalled peptide to be cleared from the ribosome by peptidyl-tRNA hydrolase (PTH). We show that *Bacillus subtilis* can survive without *trans*-translation and BrfA (Bacillus ArfA homolog), due to the presence of RqcH. We also show that expression of RqcH and its helper protein RqcP rescues the synthetic lethality of Δ*ssrA*Δ*arfA* in *Escherichia coli*. These results suggest that non-stop ribosome complexes can be disassembled and then cleared because of the tagging activity of RqcH, and that this process is essential in the absence of non-stop ribosome rescue pathways. Moreover, we surveyed the conservation of ribosome rescue pathways in >14,000 bacterial genomes. Our analysis reveals a broad distribution of non-stop rescue pathways, especially *trans*-translation and RqcH, and a strong co-occurrence between the ribosome splitting factor MutS2 and RqcH. Altogether, our results support a role for RqcH in non-stop ribosome rescue and provide a broad survey of ribosome rescue pathways in diverse bacterial species.

**Importance:** Ribosome stalling on damaged mRNA is a major problem in bacteria. It is estimated that 2-4% of all translation reactions terminate with the ribosome stalled on a damaged mRNA lacking a stop codon. Mechanisms that rescue these ribosomes, such as *trans*-translation, are often essential for viability. We investigated the functional overlap between RqcH and the non-stop ribosome rescue systems (ArfA and *trans*-translation) that are present in both *E. coli* and *B. subtilis*. Since these two species are extremely distant relatives, our work is likely to have wider implications for understanding ribosome rescue in bacteria. Furthermore, we used a bioinformatics approach to examine the conservation and overlap of various ribosome rescue systems in >14,000 species throughout the bacterial domain. These results provide key insights into ribosome rescue in diverse phyla.

## Introduction

Truncated mRNAs that lack a stop codon (non-stop mRNAs) are a major problem in bacteria (1, 2). The release factors (RF1 and RF2) require stop codon recognition to terminate translation (3, 4). Therefore, if an mRNA lacks a stop codon, the ribosome becomes stalled at the 3’ end of the message (5). Non-stop mRNAs arise from premature transcription termination, mRNA degradation by nucleases, and stop codon readthrough. Bacteria have evolved ways to rescue ribosomes from non-stop mRNAs. The primary rescue system found in nearly all bacteria, *trans*-translation, is mediated by transfer-messenger RNA (tmRNA) encoded by the *ssrA* gene, and its partner protein, small protein B (SmpB) (5, 6). SmpB bound to tmRNA senses the empty mRNA channel of the non-stop ribosome and translation then resumes on the mRNA like-domain of tmRNA (7, 8). A degradation tag, encoded by the short reading frame of tmRNA, is appended to the nascent peptide that will target the truncated protein for degradation by proteases (5, 9, 10). A stop codon at the end of the tmRNA reading frame recruits release factors for translation termination. *trans*-Translation is essential in many bacterial species, including *Mycobacterium tuberculosis*, *Shigella flexneri*, and *Neisseria gonorrhoeae* (11–13).

An alternative non-stop rescue factor, ArfA, was uncovered from a synthetic lethal screen in *E.coli* lacking *ssrA* (14). ArfA recognizes the empty mRNA channel and recruits RF1 or RF2 to terminate translation independent of a stop codon (15–17). A second alternative rescue factor, ArfB, is essential in *Caulobacter crescentus* in the absence of *trans*-translation (18). *C. crescentus* does not encode ArfA. And, although *E. coli* does encode ArfB, it is poorly expressed and not sufficient to rescue Δ*ssrA*Δ*arfA* (19). The essentiality of *trans*-translation in many bacteria and the conditional essentiality of ArfA and ArfB in the absence of *trans*-translation indicate that rescuing ribosomes from truncated mRNAs is essential (20).

*Bacillus subtilis* can survive without *trans*-translation, and encodes an alternative rescue factor called *Bacillus* ribosome rescue factor A (BrfA) (21). BrfA is homologous to ArfA, sharing 21.54% protein sequence identity when aligned with Clustal Omega (22). Like ArfA, BrfA recognizes non-stop ribosomes and recruits RF2 to terminate translation (21). BrfA is also negatively regulated by *trans*-translation, since the *brfA* transcript contains an RNase III cleavage site upstream of the stop codon and is therefore translated from a non-stop mRNA (21).

In addition to *trans*-translation and BrfA, *B. subtilis* also encodes the <u>R</u>ibosome <u>q</u>uality <u>c</u>ontrol <u>H</u>omolog RqcH. RqcH is broadly distributed in bacteria but is absent in most Alpha-, Beta-, and Gamma-proteobacteria including *E. coli* (23). RqcH acts on 50S ribosomal subunits obstructed with peptidyl-RNA. If the 70S ribosome dissociates from an mRNA and separates into 30S and 50S subunits before translation termination, the 50S subunit remains obstructed with the P-site tRNA still covalently bound to the nascent polypeptide in the exit tunnel. When this occurs, RqcH recruits alanine-charged tRNAs to the obstructed large subunit, allowing the large subunit to catalyze addition of an alanine tag to the stalled peptide in a template-independent manner. The alanine tail serves as a degron tag to target the stalled peptide for degradation (24, 25). The alanine tail also exposes the amino-acyl bond between the peptide and the tRNA to the cytoplasm, where the enzyme PTH can then hydrolyze this bond and thus free the nascent chain from the tRNA (26). The nascent peptide then diffuses, making the large subunit available for another round of translation.

Here we show that *B. subtilis* can survive without *trans*-translation and the alternative rescue factor BrfA because it encodes RqcH. We further show that RqcH and the helper protein RqcP (27) from *B. subtilis* are sufficient to rescue deletion of Δ*arfA*Δ*ssrA* in *E. coli*. Finally, we report a bioinformatic analysis of >14,000 species across the bacterial domain to determine the prevalence and distribution of *trans*-translation, ArfA/BrfA, ArfB, and RqcH. We report a strong co-occurrence between RqcH and the ribosome splitting factor MutS2. Our findings provide insight into the conservation and functional interaction between ribosome rescue systems.

## Results

### Δ*smpB*Δ*rqcH* exhibits a severe growth defect similar to Δ*smpB*Δ*brfA* in *Bacillus subtilis*

Ribosomes stall on truncated mRNAs that lack a stop codon (non-stop mRNA) since translation termination by release factors requires the presence of a stop codon in the ribosomal A-site. *B. subtilis* can rescue non-stop ribosomes using either *trans*-translation (encoded by *ssrA* and *smpB*) or BrfA (encoded by *brfA*). Deleting *brfA* and *smpB* causes a severe synthetic growth defect (Fig. 1), consistent with previously published results (21). The maximum growth rate of wild-type cells was 1.30±0.14 hour^-1^ whereas the maximal growth rate of Δ*smpB*Δ*brfA* was 0.88±0.4 hour^-1^ at 37°C. Δ*smpB*Δ*brfA* cells also exhibited a longer lag phase than wild-type cells (2.50±0.54 hours and 0.94±0.13 hours, respectively) when transferred from an overnight culture into fresh media. *B. subtilis* encodes the ribosome quality control factor RqcH. In contrast to *trans*-translation and BrfA, RqcH does not act directly on ribosomes stalled on non-stop mRNAs but adds alanine residues to the stalled peptide which is subsequently hydrolyzed from the tRNA by PTH (23, 26). Single deletion of *rqcH* had no measurable effect on growth in LB at 37°C (Fig. 1). However, the Δ*smpB*Δ*rqcH* strain exhibited a severe growth defect with a maximum growth rate of 0.64±0.16 hour^-1^ at 37°C. Δ*smpB*Δ*rqcH* also exhibited an increased lag time compared to wild-type cells. Wild-type cells took 0.94±0.13 hours to exit lag phase, whereas Δ*smpB*Δ*rqcH* took 3.94±0.75 hours to exit lag phase. When comparing Δ*smpB*Δ*rqcH* to Δ*smpB*Δ*brfA*, the Δ*smpB*Δ*rqcH* strain exhibited a significantly slower maximal growth (p = 0.0241) and significantly longer lag phase (p =0.0233). These data suggest that RqcH plays a significant role in ribosome rescue in the absence of *trans*-translation in *B. subtilis*.

**Figure 1.**
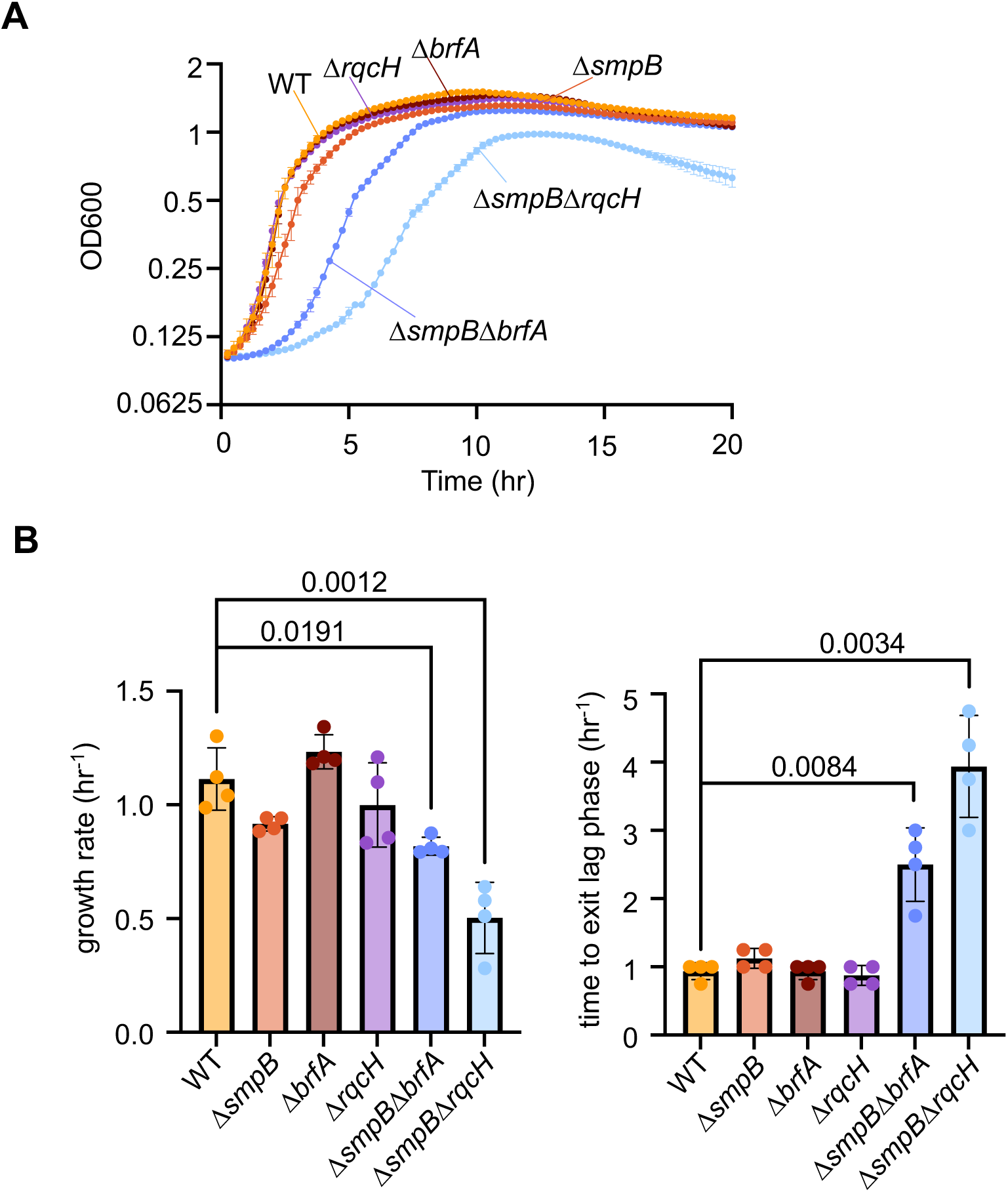
Growth of wild-type *B. subtilis,* Δ*smpB*, Δ*brfA*, Δ*rqcH::kan^R^*, Δ*smpB*Δ*brfA::kan^R^*, and Δ*smpB*Δ*rqcH::kan^R^* cells. (A) Δ*smpB*Δ*brfA::kan* and Δ*smpB*Δ*rqcH* cells exhibit a more severe growth defect and longer lag-phase under aerobic growth in LB at 37°C. (B) Bar graphs show maximal growth rates and time for cells to exit lag phase (time at which OD exceeds 0.125). Δ*smpB*Δ*rqcH* cells have an increased lag phase and slower maximal growth rate than wild type. Error bars represent the standard deviation of four independent replicates performed on different days. p-values represent results of an unpaired t-test with Welch’s correction.

### SmpB is required for cell growth in the absence of *brfA* and *rqcH* in *B. subtilis*

Deleting *smpB* and *brfA* from *B. subtilis* causes a severe, but non-lethal growth defect (Fig. 1). In contrast, deleting *smpB* and *arfA* from *E. coli* is lethal (14). Therefore, we hypothesized that *B. subtilis* can survive deletion of *smpB* and *brfA* because it encodes RqcH, which is absent in *E. coli*, and that deletion of all three factors (*smpB*, *brfA* and *rqcH*) would be lethal to *B. subtilis*. To test this, we used CRISPR interference (CRISPRi) (28, 29) to deplete SmpB from cells lacking *rqcH* and/or *brfA*. We expressed nuclease deficient Cas9 (dCas9) under the control of a xylose inducible promoter while constitutively expressing a guide RNA targeting *smpB* (sgRNA*^smpB^*). Knocking down *smpB* from Δ*brfA* or Δ*rqcH* cells decreased colony size, consistent with the growth defect observed in liquid culture, but was not lethal (Fig. 2). In contrast, when *smpB* expression was blocked in the Δ*rqcHΔbrfA* strain, colonies failed to form on plates at both 37°C and 30°C (Fig. 2B), indicating that *trans*-translation becomes essential in the absence of BrfA and RqcH.

**Figure 2.**
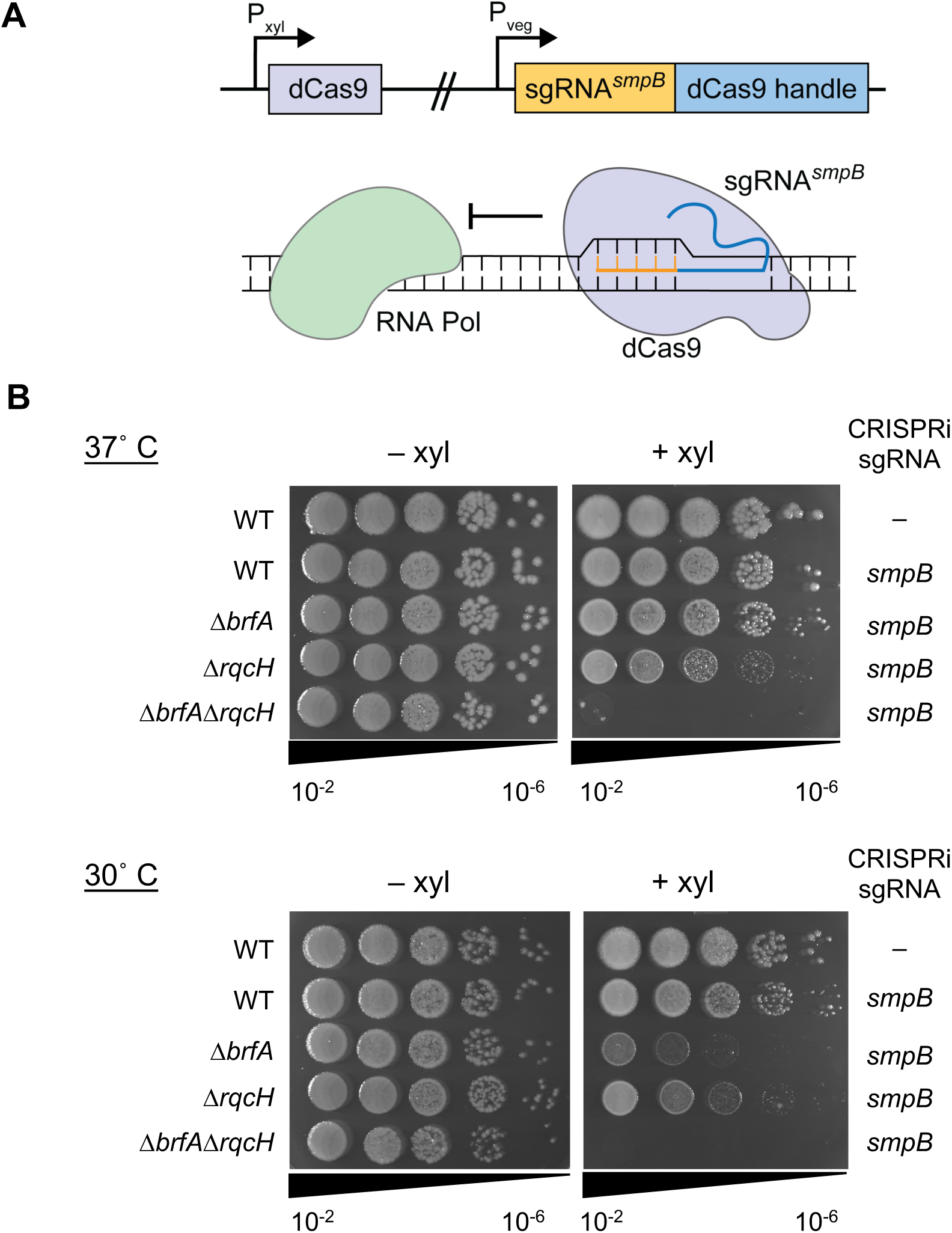
Impact of *smpB* depletion from cells lacking *brfA & rqcH.* (A) Schematic showing depletion of *smpB* using CRISPRi. Guide RNA targeting *smpB* (sgRNA*^smpB^*) under the vegetative promoter in *B. subtilis* is co-expressed with deactivated Cas9 (*dCas9*) under the control of a xylose inducible promoter resulting in transcriptional repression. (B) Strains harboring sgRNA*^smpB^* and *dCas9* were serially diluted and spot plated on LB agar with or without 1% xylose at 37°C and 30°C. SmpB depletion from Δ*brfA*Δ*rqcH* is more severe than depletion of SmpB from the single deletions. SmpB depletion from the Δ*brfA* or Δ*rqcH* single deletions is more severe at 30°C than at 37°C. Images are representative of three independent experiments.

### *B. subtilis* RqcH and RqcP rescues the synthetic lethal phenotype of Δ*ssrAΔarfA* in *E. coli*

*E. coli* lacks the RQC system but can survive deletion of *trans-*translation because of the alternative non-stop ribosome rescue system, ArfA (14). To determine whether *E. coli* lacking *ssrA* and *arfA* could survive when expressing components of the *B. subtilis* RQC pathway, we transformed MG1655Δ*ssrA::cat^R^* with a plasmid encoding *rqcH* and the helper protein *rqcP* (from *B. subtilis)* under the control of an arabinose inducible promoter. We then used phage lysate generated from MG1655Δ*arfA::kan^R^* to transduce the *ΔarfA::kan^R^*deletion into MG1655Δ*ssrA::cat^R^ E. coli* strains, harboring either empty vector or vector encoding RqcH and RqcP (pRqcH-RqcP). No colonies were recovered when Δ*arfA::kan^R^* was transduced into the Δ*ssrA* strain containing empty vector. In contrast, colonies were recovered on LB with 1% arabinose when Δ*arfA::kan^R^* was transduced into the Δ*ssrA* strain containing plasmid encoding arabinose-inducible RqcH and RqcP. These cells did not grow on plates in the absence of arabinose, indicating that expression of RqcH/RqcP is required for survival in this condition (Fig 3A). In liquid culture, Δ*arfA*Δ*ssrA* containing pRqcH-RqcP grew better in the presence of arabinose (Fig 3B). These results demonstrate that RqcH and RqcP are sufficient to support *E. coli* survival when *trans-*translation and ArfA are absent.

**Figure 3.**
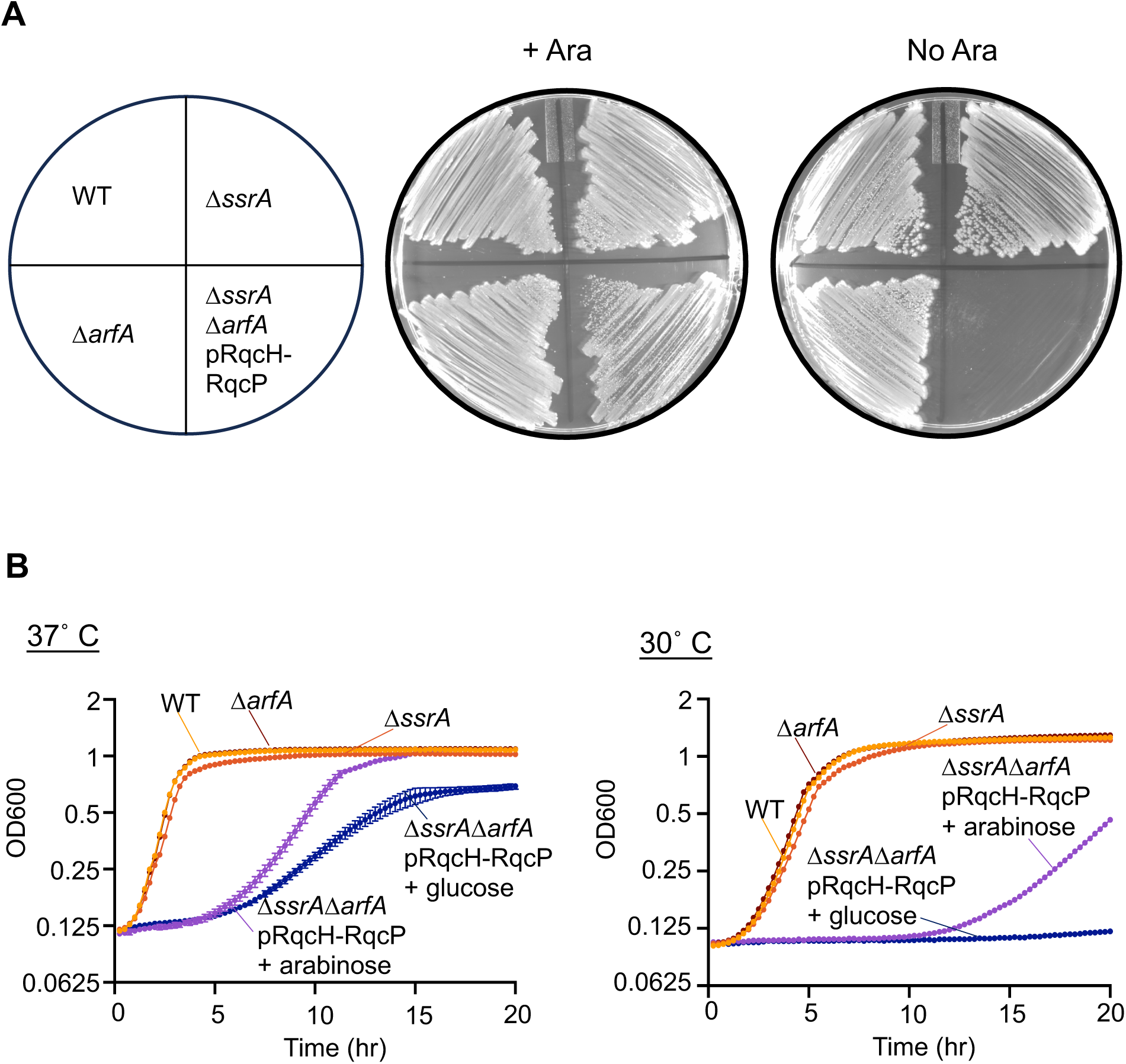
*E.coli* can survive the synthetic lethality of Δ*ssrA::cat^R^*Δ*arfA::kan^R^*when provided with *B. subtilis rqcH and rqcP*. (A) Strains depicted in the schematic were grown on LB plates with or without 1% arabinose to induce expression of RqcH/RqcP. (B) Growth curves in LB of *E. coli* Δ*arfA*::*kan^R^*, Δ*ssrA*::*cat^R^*and Δ*ssrA::cat^R^*Δ*arfA::kan^R^* strain harboring the plasmid expressing RqcH and RqcP. Δ*ssrA*Δ*arfA* with pRqcH-RqcP grows better with arabinose induction of RqcH and RqcP. However, cells are not recovered to the wild type growth rate. Error bars represent the standard deviation of cells grown from independent colonies in triplicate.

### Ribosome rescue pathways are broadly conserved in bacteria

*B. subtilis* RqcH/RqcP rescued the synthetic lethality of Δ*ssrA*Δ*arfA* in *E. coli*, a species which lacks RqcH. To investigate the conservation of various ribosome rescue pathways in bacteria, we surveyed 14,479 representative reference genomes for the presence of genes encoding tmRNA, SmpB, ArfA, ArfB, and RqcH. Consistent with previous reports (30), we detected the genes encoding *trans*-translation in most bacterial genomes (Fig. 4) (Table 1). *smpB* was detected in 95.3% of the representative genomes surveyed and *ssrA* was detected in 94.2%. 95.8% of genomes contained either *smpB* or *ssrA* (Fig. 4B) (Table 1). Genomes that lacked *ssrA* were found mainly in Mycoplasmatota and Bacteria incertae sedis. However, most of these genomes still encoded *smpB*, indicating *ssrA* may still be present, but was not identified in our search. *arfB* is widely distributed across the bacterial domain and was conserved in 54.7% of surveyed genomes. Most Gram-positive phyla, including Firmicutes, lack *arfB*. *arfA* was the least conserved rescue pathway, and is restricted specifically to Gammaproteobacteria, some Betaproteobacteria, and 10 species in the *Bacillus subtilis* group. *rqcH* was identified in 23.6% of the surveyed genomes and was found in phyla across the bacterial domain, except Proteobacteria. The broad conservation of RqcH and the presence of an RqcH homolog in eukaryotes are consistent with RqcH being present in the last universal common ancestor (31).

**Figure 4.**
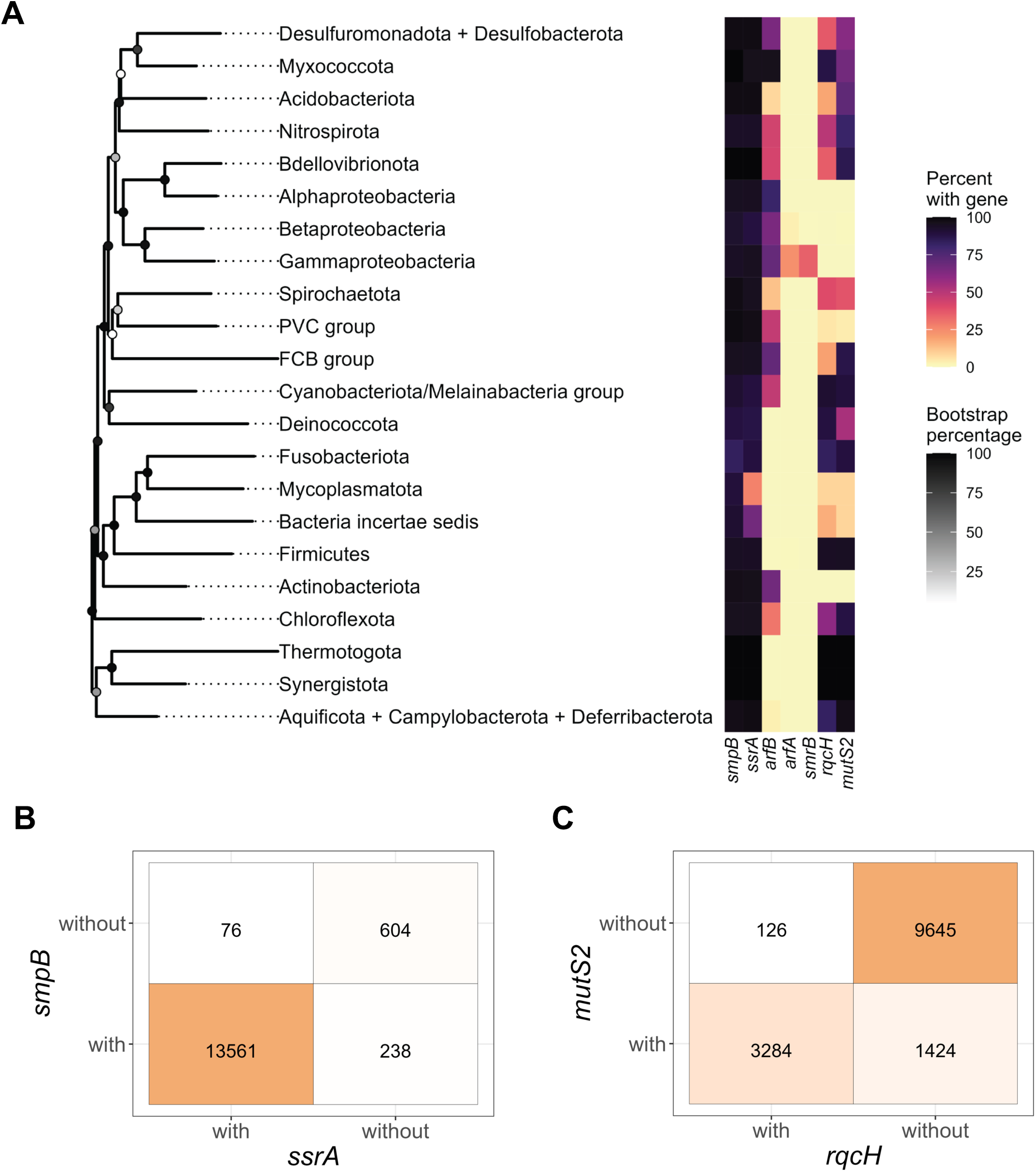
Conservation of ribosome rescue pathways across >14,000 bacterial genomes. *rqcH* and *mutS2* co-occur across bacterial phyla. (A) Heatmap showing the percent of genomes per phyla containing *smpB*, *ssrA*, *arfA*, *arfB*, *smrB*, *rqcH*, and *mutS2*. The tree was built using 16S rRNA genes from a single representative species for each phylum. NCBI accession numbers for the representatives are listed in Table 1. Bootstrap percentages are shown as darkened circles at each node. (B) Co-occurrence matrix of *smpB* and *ssrA*. The values represent the number of genomes with or without these genes. (C) Co-occurrence matrix of *rqcH* and *mutS2*. The values represent the number of genomes with or without these genes.

**Table 1:**
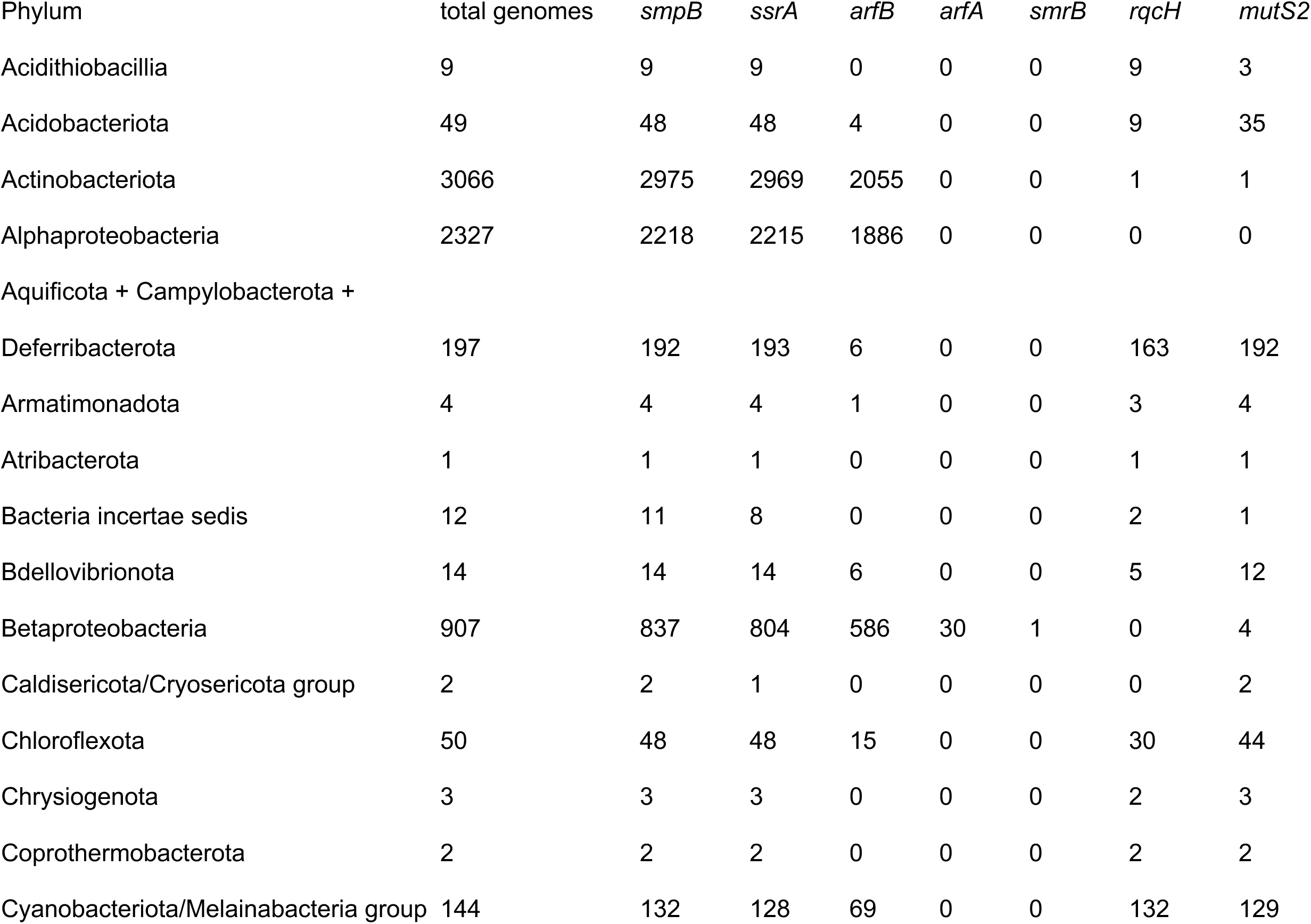
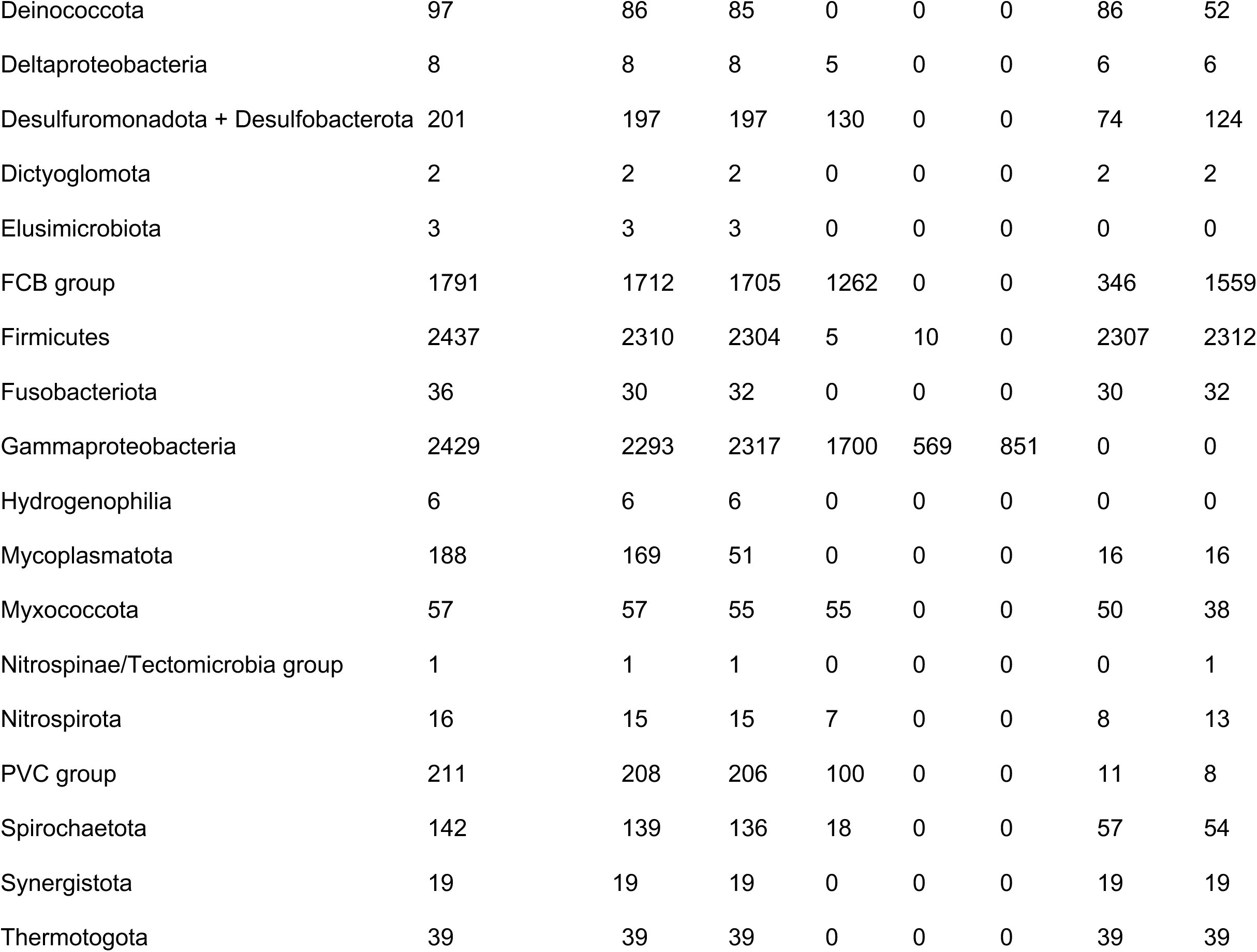
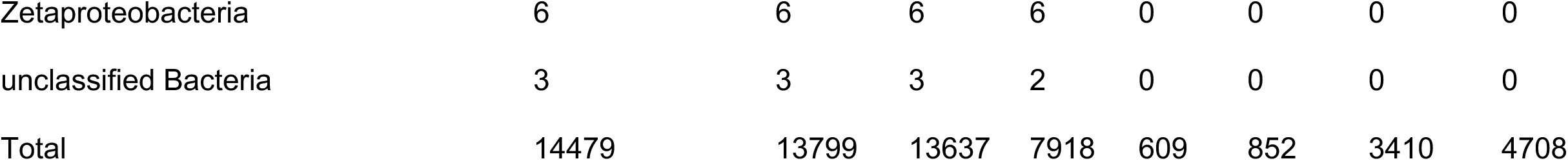
Conservation of ribosome rescue pathways across 14,479 genomes.

### RqcH and MutS2 frequently co-occur in bacterial genomes

Two newly discovered ribosome quality control proteins are SmrB and MutS2. Both proteins contain SMR domains and bind collided ribosomes (31–33). MutS2 splits stalled ribosomes from the mRNA, leaving an obstructed large ribosomal subunit that is a substrate for RqcH (31). MutS2 and RqcH are both found throughout the bacterial domain and have homologs in eukarya (31). Since MutS2 generates substrates for RqcH, we investigated how frequently MutS2 and RqcH co-occur in bacterial genomes. 96% of genomes that encoded RqcH also encoded MutS2 (Fig. 4C). Of the genomes that have lost RqcH, 87% have also lost MutS2. These data indicate that there is a high degree of co-occurrence between RqcH and MutS2.

### RqcH ribosome rescue activity is decreased, but not abolished, in the absence of MutS2

Our bioinformatic analysis revealed a strong co-occurrence between RqcH and MutS2 in bacterial genomes, suggesting that RqcH is dependent on the ribosome splitting activity of MutS2. To test this, we compared the impact of SmpB depletion from Δ*brfA*Δ*rqcH* cells to SmpB depletion from Δ*brfA*Δ*mutS2* cells. In the Δ*brfA* background, when SmpB is depleted, cells are totally reliant on RqcH. In three independent experiments, SmpB depletion from Δ*brfA*Δ*mutS2* exhibited a more severe fitness defect than SmpB depletion from the Δ*brfA* single deletion, indicating that MutS2 is a major splitting factor acting upstream of RqcH (Fig. 5). However, this fitness defect was not as severe as SmpB depletion from Δ*brfA*Δ*rqcH*. These data indicate that RqcH retains some function in the absence of MutS2, and that there may be additional factors capable of splitting ribosomes from nonstop messages prior to rescue by RqcH.

**Figure 5.**
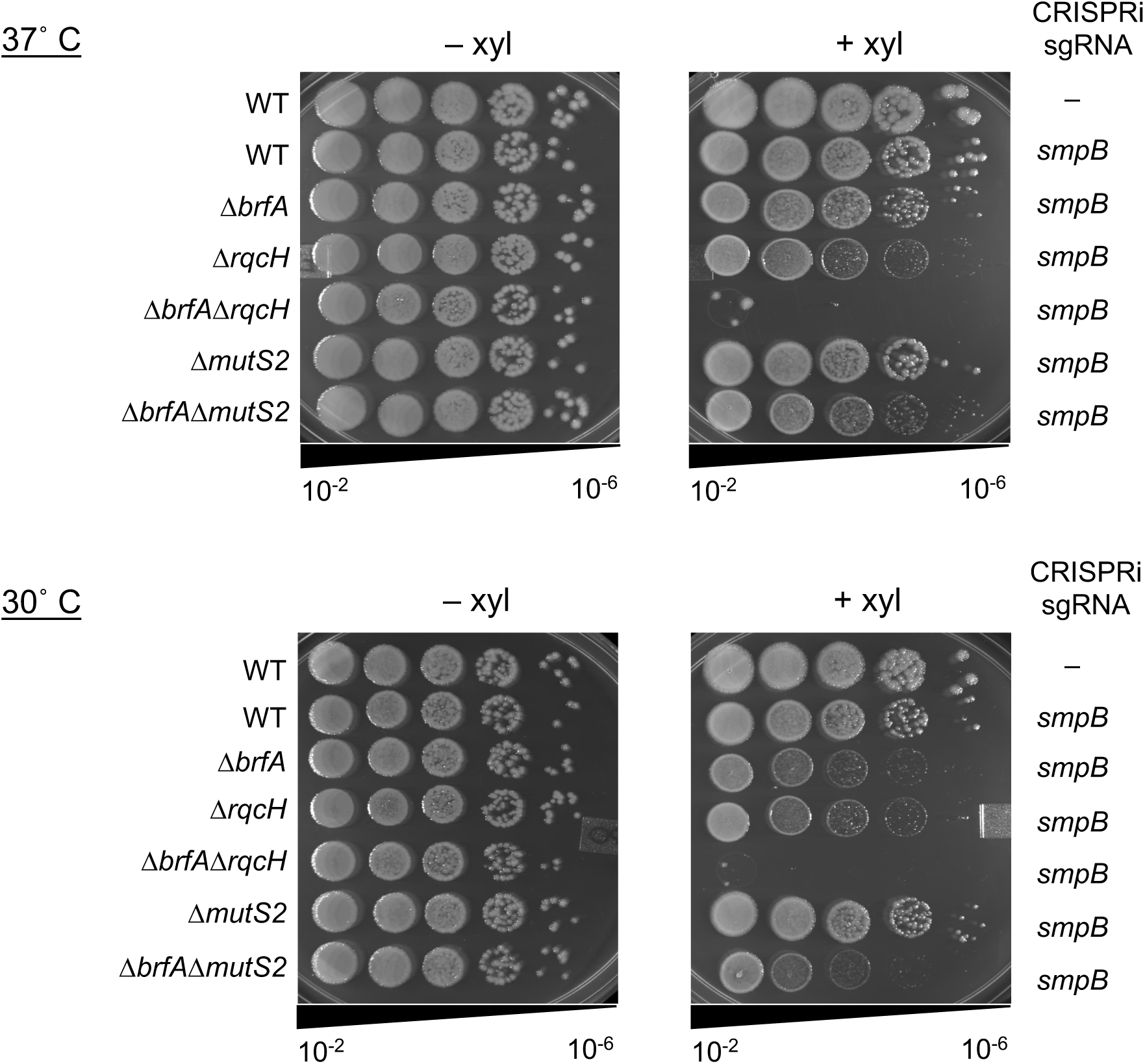
The RQC pathway can still function in the absence of MutS2. Strains harboring sgRNA*^smpB^* and *dCas9* were serially diluted and spot plated on LB agar with or without 1% xylose at 37°C and 30°C. Depleting SmpB from Δ*rqcH* cells resulted in a severe growth and survival defect at both 30°C and 37°C. In contrast, depleting SmpB from Δ*mutS2* cells results in a modest growth defect in wild-type cells at both 37°C and 30°C. Similarly, depleting SmpB from Δ*brfA*Δ*mutS2*, resulted in growth and survival defects that were more severe than depleting SmpB from Δ*brfA*, but not as severe as depleting SmpB from Δ*brfA*Δ*rqcH*, indicating that RqcH is not absolutely dependent on the activity of MutS2. Image shows a representative from three independent experiments.

## Discussion

Previous work demonstrates that bacteria can survive without *trans*-translation only if they encode one or more alternative non-stop rescue factors, either ArfA or ArfB (12, 14, 18). However, these studies were restricted mainly to Proteobacteria, which lack the recently discovered ribosome rescue factor RqcH. RqcH is highly conserved in bacterial phyla outside of Proteobacteria (Fig. 4). Therefore, we investigated the essentiality of the non-stop rescue factors in *B. subtilis*, a model Firmicute that encodes RqcH. We found that *B. subtilis* lacking all the canonical non-stop rescue factors (*trans*-translation, ArfA and ArfB) can survive because it encodes RqcH (Fig. 1 and 2). Moreover, expression of RqcH and its helper protein RqcP in *E. coli* rescued the well-documented synthetic lethality of Δ*ssrA*Δ*arfA* in this species (Fig. 3). These results suggest that RqcH/RqcP can support viability in the absence of *trans*-translation and the alternative non-stop ribosome rescue pathways.

RqcH does not act directly on non-stop ribosomes but can still support cell survival in the absence of *trans*-translation and *arfA* in both *B. subtilis* and *E. coli*. These results suggest that ribosomes can dissociate from non-stop messages and subsequently be rescued by the activity of RqcH and PTH. How do ribosomes dissociate from non-stop mRNAs? Recently, MutS2 was identified as a ribosome splitting factor that can remove collided ribosomes from mRNAs (31). Since MutS2 generates substrates for RqcH, we determined how frequently MutS2 and RqcH co-occur in bacterial genomes. We found an extremely high co-occurrence of MutS2 and RqcH in genomes across the bacterial domain (Fig. 4). However, experimentally, we found that RqcH does not have a strict requirement for MutS2 (Fig. 5). SmpB depletion from Δ*brfA*Δ*mutS2* cells exhibited decreased fitness, but not as severe as depletion from Δ*brfA*Δ*rqcH* cells (Fig. 5). Moreover, RqcH/RqcP were sufficient to rescue viability of Δ*arfA*Δ*ssrA* in *E. coli*, a species that does not encode MutS2 (Fig. 3). Therefore, MutS2 is unlikely to be the only ribosome splitter. One potential ribosome splitter is YbnA/HflX, a universally conserved GTP-binding protein that functions as a ribosome splitting factor and is important during stressful conditions such as heat shock where ribosome stalling may increase (34). However, HflX is only known to act on hibernating ribosomes or on ribosomes carrying a deacylated tRNA in the P-site (35, 36).

Neither RqcH or MutS2 were detected in the Proteobacteria we surveyed (Table 1). Gamma-proteobacteria encode SmrB, an SMR domain-containing protein which binds to collided ribosomes but does not split them (33). Instead, SmrB possesses endonuclease activity that cleaves mRNA between collided ribosomes. *trans*-Translation, ArfA, and ArfB all require truncated mRNA and their rescue activity is undetectable on ribosomes that are stalled mid-message (37–40). Therefore, in the absence of RqcH, cells are more reliant on *trans*-translation and ArfA or ArfB to rescue ribosomes, and it becomes more important for mRNA cleavage to occur. Thus, loss of RqcH could be the pressure that selected for the nuclease activity of SmrB.

Although RqcH can support viability in the absence of *trans*-translation and is conserved in 24% of genomes, the genes encoding *trans*-translation are present in nearly all bacterial genomes (>95%) (Table 1). Even obligate intracellular bacteria with reduced genomes such as *Rickettsia rickettsii*, *Chlamydia trachomatis*, *Coxiella burnetii,* and *Buchnera aphidicola* retain *trans*-translation. *trans*-Translation provides a means to both rescue the stalled ribosome and target the stalled peptide for degradation by proteases in one pathway. In contrast, the RQC pathway requires a ribosome splitter, addition of a degron tag to the stalled peptide by RqcH, and the activity of PTH to hydrolyze the stalled peptidyl-tRNA and clear the obstructed ribosome. Therefore, *trans*-translation provides the most efficient mechanism to rescue non-stop ribosomes, which may explain its near-total conservation.

While the mRNA cleavage activity of SmrB allows *trans*-translation to be the dominant ribosome rescue pathway in *E. coli*, *B. subtilis* has evolved to also rely heavily on RqcH. It is notable that the growth defect of Δ*smpB*Δ*brfA* in *B. subtilis* exhibits similar severity to Δ*smpB*Δ*rqcH*. This finding indicates that obstructed 50S ribosomal subunits are a major problem in *B. subtilis*. Future work is needed to identify the major sources of these obstructed subunits and to identify factors that split stalled ribosomes in *B. subtilis*.

## Materials and Methods

### Strains and media

Strains were derived from *Bacillus subtilis* 168 *trpC2* and grown in LB media (10g tryptone, 5g yeast extract, 5g NaCl per liter) at 30°C or 37°C with aeration where indicated (Table 1). Gene deletions were made by transforming genomic DNA from the BKK collection (41) into the lab’s naturally competent wild type. The pDR244 marker loop-out plasmid was used to create a marker less *smpB* deletion from the BKE collection and a marker less *brfA* from the BKK collection. Final concentrations of antibiotics used for selection included 100ug/mL ampicillin, 20ug/mL kanamycin, 100ug/mL spectinomycin, 5ug/mL chloramphenicol, and 1X MLS (1ug/mL erythromycin and 25ug/mL lincomycin).

### Growth curves for *B. subtilis*

Cells were grown overnight in LB with aeration and back diluted to a starting OD600 of 0.5 and serially diluted to OD600 of 0.05, 0.005, and 0.0005. The cells were grown for 24h at 37°C, shaking at 2mm amplitude using Thermo Scientific 96-well flat bottom plates (Cat.No. 167008). OD600 values were obtained in 15min intervals from the BioTek Synergy H1 microplate reader, Gen5 3.11. Growth rates at OD600=0.05 were measured using non-linear regression of logistic growth and statistical analysis was performed using an unpaired t-test followed by a Welch’s correction using GraphPad Prism version 10.1.1 for macOS.

### Plasmid construction

RqcH from *Bacillus subtilis* was amplified using primers KC53 & KC54 and gel extracted. pBAD322 digested with EcoRI/NcoI was used as the plasmid backbone for Gibson assembly (42) of the PCR product resulting in pKC353. IDT gblock of RqcP with homology to pKC353 was then Gibson assembled after pKC353 digestion with NcoI/XbaI to create pKC426 (Genbank ID). Plasmids were transformed into Dh5alpha cells and selected on ampicillin plates containing 1% arabinose. Whole Plasmid Sequencing was performed by Plasmidsaurus using Oxford Nanopore Technology with custom analysis and annotation.

### CRISPR interference

To create the sgRNA targeting SmpB, we amplified pJMP2 using primer sp60 (29) and reverse primer HRH175 (43). The Phusion PCR product was DpN-1 treated for 2.5hrs at 37°C followed by PNK treatment for 1h. The resulting band was gel extracted and ligated overnight at room temperature using T4 DNA ligase (NEB). pKC349 was transformed into Dh5alpha and selected on ampicillin and sequencing was performed by Plasmidsaurus. Genomic DNA of *B. subtilis* harboring dCas9 under an inducible xylose promoter at *lacA* (43) was transformed into Δ*rqcH::kan*, Δ*brfA,* and Δ*brfA*Δ*rqcH::kan* cells. pKC349 was ScaI digested and transformed into the above backgrounds for integration into *amyE.* Strains were grown in LB for 4h at 37° and back diluted to a final OD600 of 0.05 in 1XTbase + 1mM MgS04. The cultures were serially diluted and 10ul were spotted onto LB agar containing 1% xylose and incubated at 30°C or 37°C overnight.

### Transduction

Phage lysate was obtained from *E. coli* Δ*arfA::kan*. MG1655Δ*ssrA::cat* was electroporated with pKC353 or pKC426 and grown on LB agar containing ampicillin. Recipient cells were transduced with phage lysate (44–49) and selected on media with and without 1% arabinose to induce expression of RqcH (pKC353) and RqcH with RqcP (pKC426).

### Growth curves for *E. coli*

Cells were grown overnight in LB and strains harboring the pRqcH/RqcP plasmids were grown with 100ug/ml ampicillin and 1% arabinose to maintain the plasmid. The overnight cultures were back diluted to a starting OD600 of 0.05. The cells were grown in the presence of 1% arabinose or 1% glucose where indicated for 24h at 30°C or 37°C, shaking at 2mm amplitude using Thermo Scientific 96-well flat bottom plates (Cat.No. 167008). OD600 values were obtained in 15min intervals from the BioTek Synergy H1 microplate reader, Gen5 3.11.

### Gene detection

Genes were detected in a database of 14,479 representative prokaryotic genomes from NCBI RefSeq (50) using HMMER v3.3 (nhmmer) (hmmer.org) with an E-value cutoff of 0.05. HMMER profiles were built using ≥ 5 genes as annotated by the NCBI Prokaryotic Genome Annotation Pipeline (50). At least one gene sequence from every taxa expected to have the gene based on a preliminary search was included. In Alphaproteobacteria like *Caulobacter crescentus*, the *ssrA* gene is interrupted by an internal loop that is excised from the final RNA product (51). Thus, two separate HMMER profiles were built and searches were performed for *ssrA* to separately probe Alphaproteobacteria and other phyla. Proteins closely resembling *B. subtilis* RqcH are described in NCBI as “NFACT RNA binding domain-containing proteins.” Only genes for proteins with this description were used when building the *rqcH* HMMER profile. The domain architecture of MutS2 is similar to that of MutS1, but the homology is only shared over approximately 1000 bp. Therefore, hits were filtered for a coverage of >1250 bp to remove erroneous *mutS1* hits. Similarly, *arfB* shares approximately 100 bp of homology to *prfB*, so hits were filtered for a coverage of >150 bp. SMR domains in other proteins resemble the C-terminus of *smrB*, so hits were filtered for a coverage of >400 bp.

### Phylogenetics

16S rRNA sequences of all genomes were identified and acquired using BLAST v2.13.0 (52), aligned using MAFFT v7.453 (53), and applied to FastTree v2.1.11 (54) to infer a maximum likelihood tree. FastTree produces unrooted phylogenies, so trees were midpoint rooted using the phangorn v2.11.1 package (55). Representatives from each phyla were randomly selected and used to subset the built 16S tree. The identities of the randomly selected genomes can be found in Table 1. Taxonomic classification was assigned to genomes using the NCBI Taxonomy database (56). Phyla were named using the conventions in Coleman et al. 2021 (57). The tree was visualized using ggtree v3.6.2 (58).

## Acknowledgements

HAF, KC and CRP were supported by NIH R35GM147049. KC was supported by a Dean’s Scholars Diversity Fellowship from Cornell University. CRP was supported by a Graduate Research Fellowship from the National Science Foundation. We thank Kenneth Keiler, Kevin England, and Daniel Tetreault for helpful feedback on the manuscript.

---

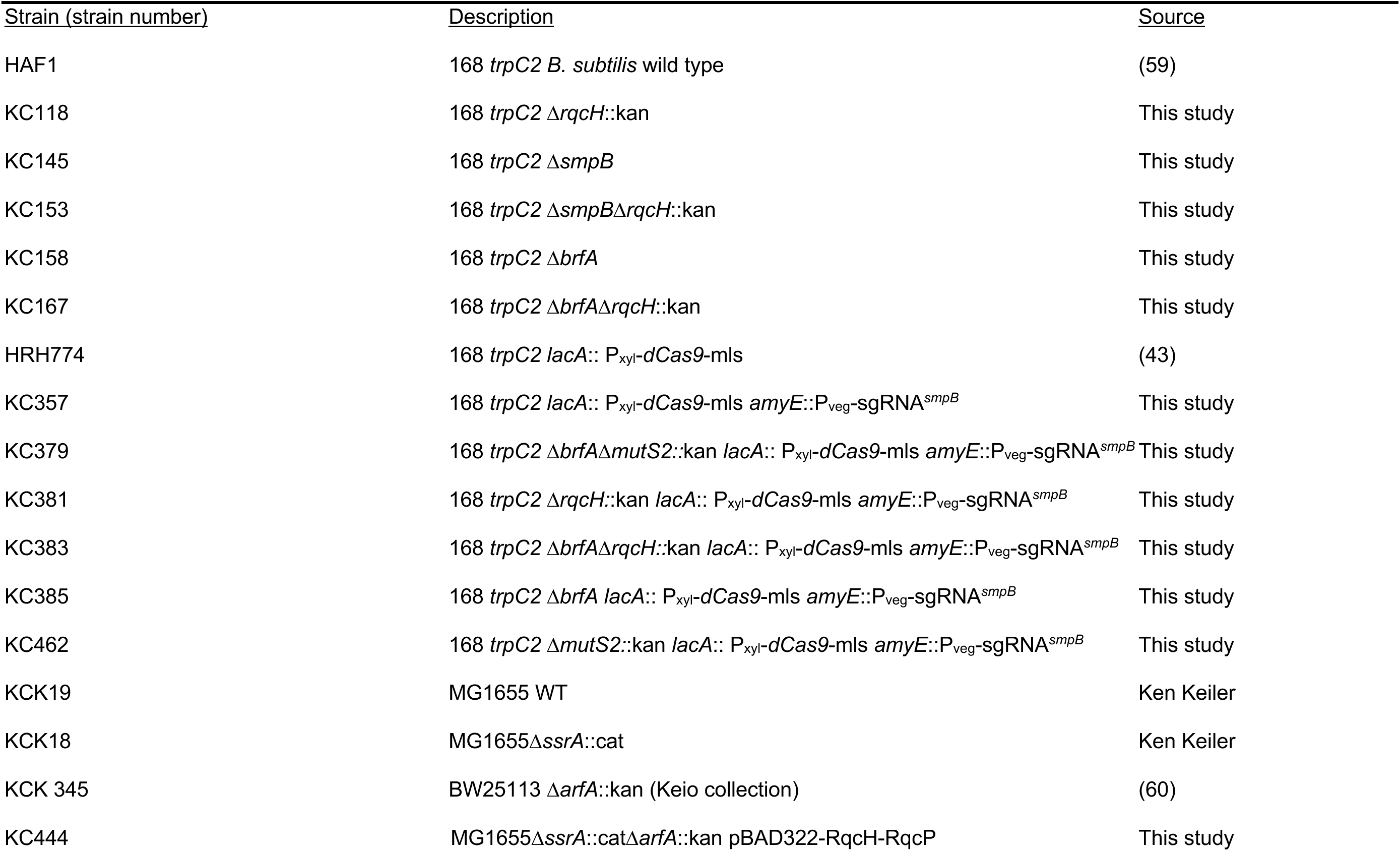
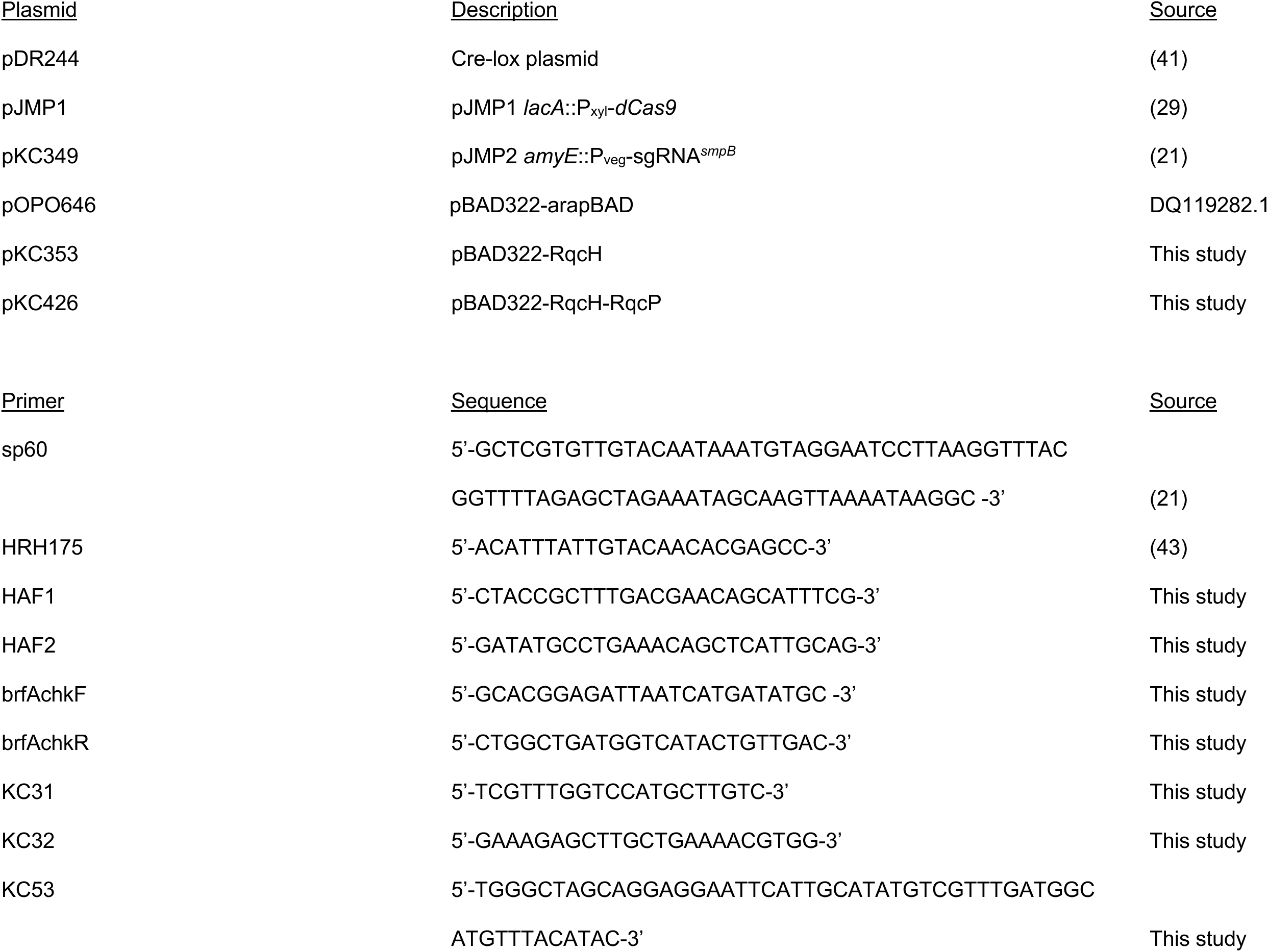
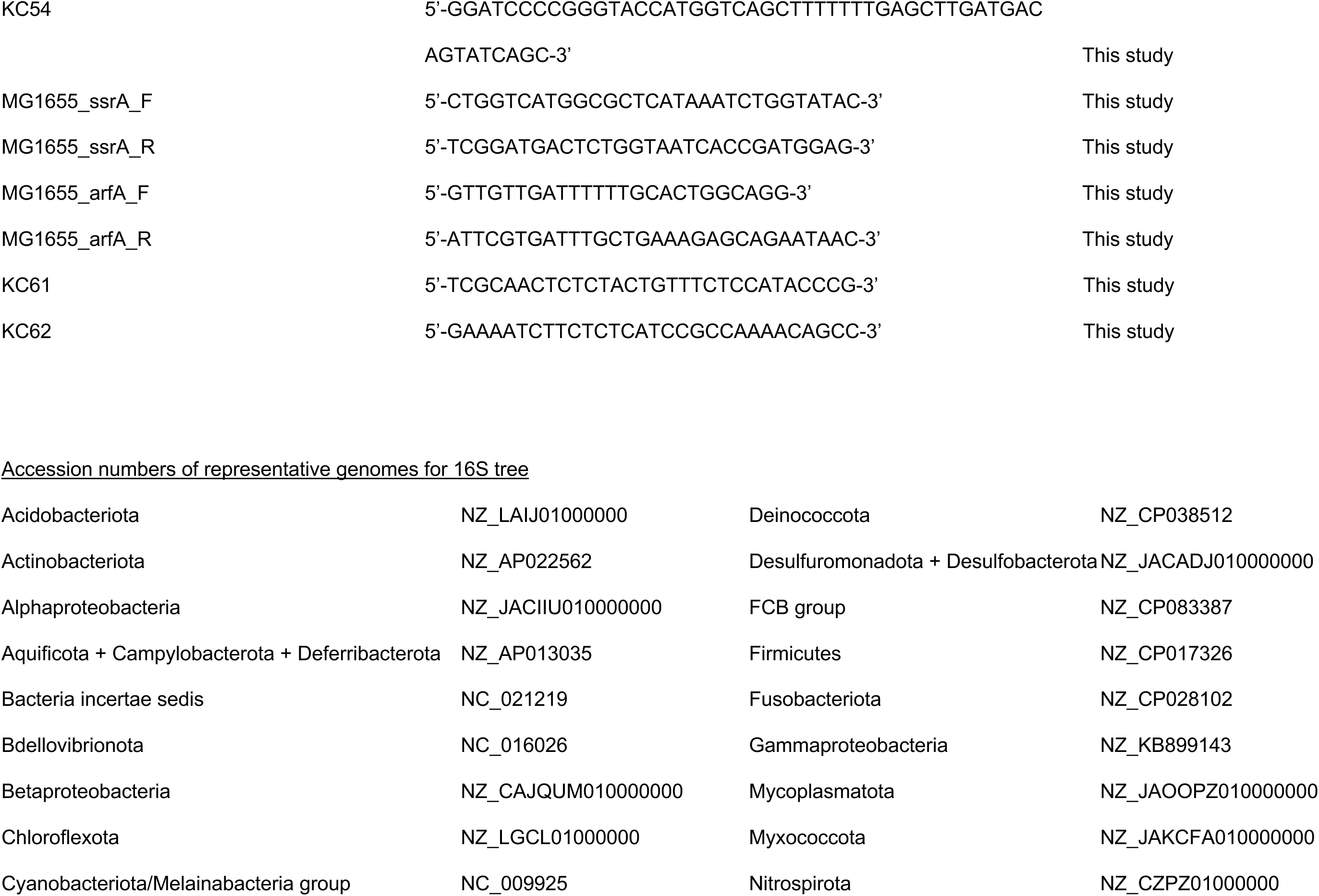
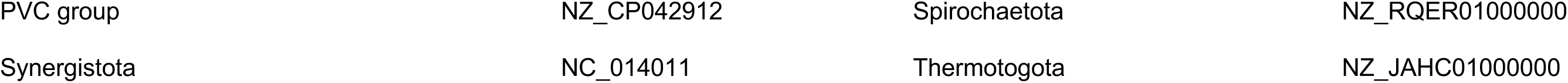

